# Gene Editing and Aging: A Bibliometric Analysis of Global Trends and Frontier Themes (2015-2024)

**DOI:** 10.1101/2025.05.01.651618

**Authors:** Lening Chen, Hui Li, Yingxia Zhu, Zhimin Zheng, Junying Wang, Hongxiong Wang, Wei Huang, Yongli Luo

**Affiliations:** Department of Geriatrics, The Third People’s Hospital of Yunnan Province, The Second Affiliated Hospital of Dali University, 292 Beijing Road, Kunming 650011, Yunnan Province, China; Library, Kunming Medical University, 1168, Chunrong West Road, Yuhua Street, Kunming 650500, Yunnan Province, China; Department of Geriatrics, General Hospital of Ningxia Medical University,804 Shengli South Street,Ningxia 750004, China

**Keywords:** Genome Editing, Aging, Bibliometric Analysis, CRISPR-Cas9, Telomere

## Abstract

**Objective:** The accumulation of DNA damage and mutations is a key contributor to aging. Recent studies have shown that disrupting the Beclin 1-BCL2 autophagy regulatory complex through gene editing can extend lifespan in mice. The precise application of gene editing technologies offers a promising strategy for aging. This study conducted a bibliometric analysis to map the knowledge landscape of gene editing and aging.

**Methods:** We retrieved publications related to genome editing and aging from the Web of Science Core Collection, covering the period from 2015 to 2024. The data were analyzed using VOSviewer and R package Bibliometrix. These tools enabled us to identify the most productive researchers, journals, institutions, countries and visualized current trends, emerging research hotspots.

**Results:** A total of 982 publications on genome editing and aging were identified. The United States (n=285) and China (n=214) form a dual-core structure leading global output. Harvard University (n=116) emerged as the most prolific institution. Scientific Reports was the top-publishing journal, with 23 articles in 2024. ZHANG Y (n=12, citations=102, H-index=6) was identified as the most productive author. KIM E’s 2017 publication in Nature Communications (TC=494, TC/year=54.9, NTC=9.33) has had a significant and ongoing impact. The analysis indicates that future directions will include CRISPR optimization and AI-assisted genomic analysis.

**Conclusion:** This study presents the first comprehensive bibliometric analysis and visualization of the knowledge structure in gene editing and aging research up to 2024. It offers researchers a detailed overview of current developments, trends, and emerging frontiers in this rapidly evolving domain.

## 1. Introduction

According to the OECD Health at a Glance 2023 report, more than 242 million people in OECD countries were aged 65 and over in 2021, including over 64 million who were at least 80 years old [1]. This demographic shift is significantly increasing the burden of age-related diseases. The human body is composed of approximately 3×10^13^ cells [2], each of which plays a specific role in maintaining life and overall functionality. Aging involves the progressive dysfunction of virtually every organ across almost all organisms. A common feature of aging is the accumulation of genetic damage over time [3]. Over the past two decades, numerous studies on the aging transcriptome have aimed to elucidate its underlying mechanisms. Renk et al. surveyed reports of age-related changes in basal gene expression across eukaryotes from yeast to humans and identified six gene expression hallmarks of cellular aging: (1) downregulation of genes encoding mitochondrial proteins; (2) downregulation of the protein synthesis machinery; (3) dysregulation of immune system genes; (4) reduction in growth factor signaling; (5) constitutive responses to stress and DNA damage; and (6) dysregulation of gene expression and mRNA processing [4]. The first evidence supporting a candidate from signature-based screening KU0063794 showed that it significantly extended both lifespan and healthspan in aged C57BL/6 mice [5]. Additional evidence comes from studies demonstrating that disruption of the Beclin 1-BCL2 autophagy regulatory complex promotes longevity in mice [6]. Cells from aged humans and model organisms accumulate somatic mutations in nuclear DNA[7], and deficiencies in DNA repair mechanisms have been shown to accelerate aging in mice and underlie several human progeroid syndromes, such as Werner syndrome, Bloom syndrome, xeroderma pigmentosum, trichothiodystrophy, Cockayne syndrome, and Seckel syndrome [8-10]. This is particularly relevant when DNA damage affects stem cells, impairing their capacity for tissue renewal or leading to their depletion. Such impairments promote aging and increase susceptibility to age-related diseases [11].usal evidence linking the lifelong accumulation of genomic damage to aging has emerged from studies in both mice and humans.

In 1977, British biochemists F. Sanger and A. R. Coulson developed the first generation of sequencing technology [12]. Over the past 15 years, gene sequencing technology has advanced rapidly, with the commercialization of high-throughput DNA sequencing methods, commonly referred to as next-generation sequencing (NGS) [13]. Currently, multi-omics approaches including transcriptomics, metabolomics, and epigenomics are being employed to assess molecular signatures associated with longevity and aging. In 2012, two research groups reported that purified Cas9, derived from Streptococcus thermophilus or Streptococcus pyogenes, can be guided by CRISPR RNAs (crRNAs) to cleave target DNA in vitro [14]. This is achieved using sequence-specific nucleases such as zinc finger nucleases (ZFNs), transcription activator-like effector nucleases (TALENs), and CRISPR/Cas9 to create double-strand breaks (DSBs) in DNA, thereby triggering cellular repair mechanisms such as non-homologous end joining (NHEJ) or homology-directed repair (HDR). These mechanisms facilitate targeted gene insertion, deletion, or replacement [14-16]. Humanity has progressed from gene sequencing to gene editing a technology that enables precise modification of DNA sequences at the genomic level. The intersection of genome editing and aging has emerged as a transformative area in biomedical research, where breakthroughs in precision editing tools are being leveraged to unravel the molecular underpinnings of aging and develop novel strategies to delay its onset. Bibliometric analysis (BA) is widely used in medical and healthcare research to evaluate the development of specific fields [17]. A bibliometric review can reveal the extent of scholarly attention devoted to a particular topic. Furthermore, by presenting and summarizing relevant studies, bibliometric analysis enables researchers to identify emerging trends in article publication, journal activity, research performance, collaboration patterns, and the intellectual structure within a given domain [18]. This study aims to analyze publication trends and construct a visualized knowledge network of research related to gene editing and aging from 2015 to 2024. Notably, the findings will provide researchers with a comprehensive overview of current knowledge, while identifying global research trends and cutting-edge themes in the field of gene editing and aging.

## 2. Methods

### 2.1. Data sources

The Web of Science Core Collection (https://www.webofscience.com/) was selected as the primary database for this study. This authoritative resource indexes over 12,000 high-impact scholarly journals across a wide range of disciplines, including the natural sciences, engineering, biomedical sciences, social sciences, and arts & humanities. A distinctive feature of the Web of Science Core Collection is its comprehensive citation indexing, which systematically tracks cited references by author, source, and publication year. This powerful citation analysis capability enables researchers to trace the historical development of scientific literature and precisely identify emerging research trends. Its widespread adoption within the academic community further underscores its credibility and utility. To identify relevant academic publications, we conducted a comprehensive literature search in the Web of Science Core Collection (WoSCC) using a targeted search strategy. The search query was as follows: TS=((Editing, Gene) OR (Genome Editing) OR (Editing, Genome) OR (Base Editing) OR (Editing, Base)) AND TS=((Biological Aging) OR (Aging, Biological) OR (Aging) OR (Senescence)) As of March 10, 2025, a total of 1,739 documents were retrieved. We then applied a series of screening criteria to enhance the precision of our analysis. These included language (English only), document type (research articles), and publication timeframe (2015-2024). After applying these filters, 982 research articles were retained for further analysis (Figure 1).

**Figure. 1.**
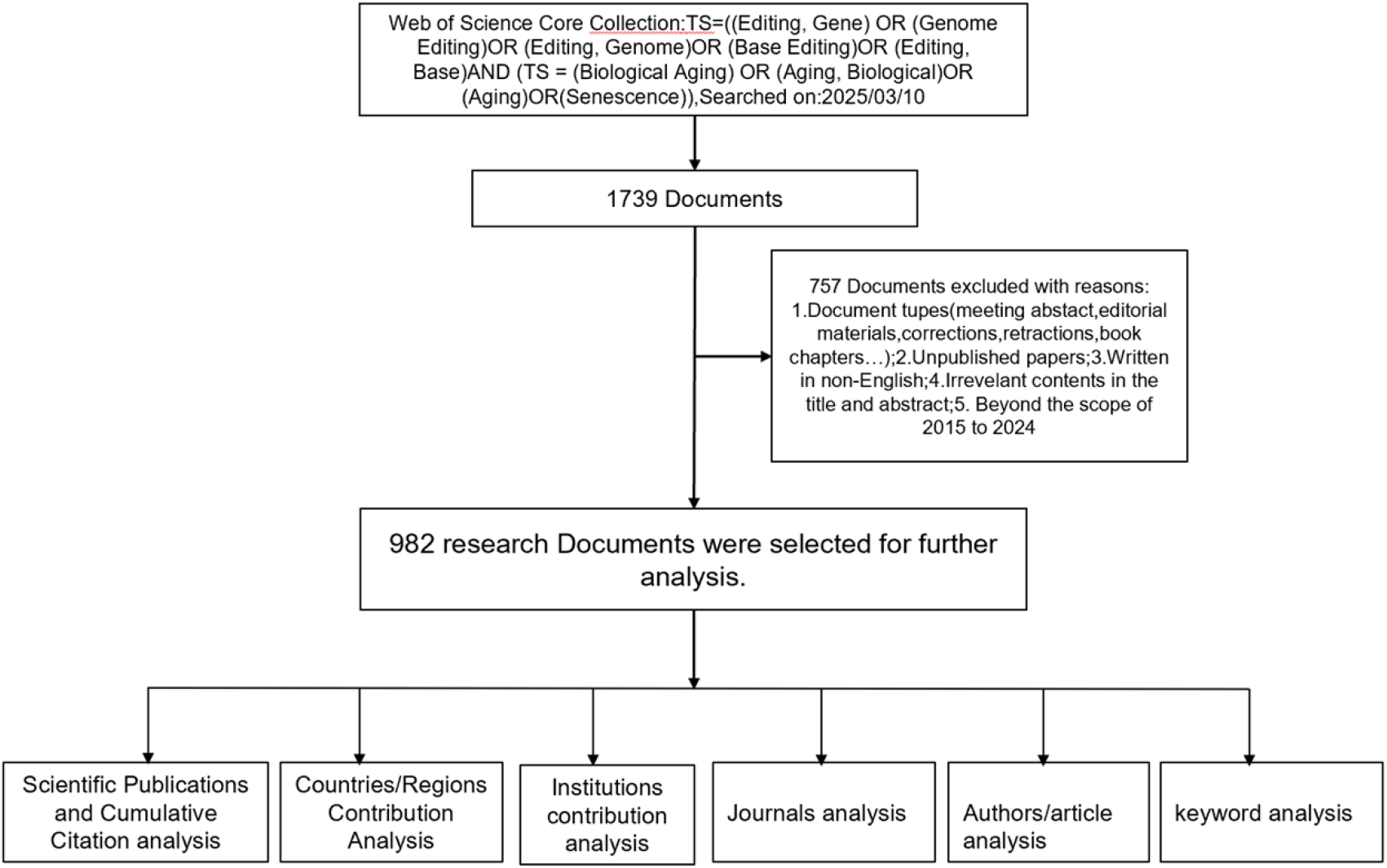
The workflow of data collection and bibliometric analysis

### 2.2. Bibliometric analysis

Bibliometrics has evolved into a comprehensive statistical approach that not only quantifies the volume of publications but also assesses emerging trends and shifts in research focus within specific subject areas. This evolution reflects its development from traditional bibliometrics to information metrics, with increasing integration into scientometrics and informetrics. Visualization is a crucial component of bibliometric analysis, as it effectively illustrates the structure and evolution of a research field.

In this study, we used VOSviewer and the Bibliometrix R package to analyze the data. After screening the records based on titles and abstracts, we downloaded the selected publications in Full Record and Cited References format, and imported the search results into R to analyze publication trends and identify major contributors. Bibliometrix and VOSviewer were then employed to construct a knowledge network, a collaboration network, a keyword co-occurrence network, and a co-citation reference network. In VOSviewer visualizations, node size is positively associated with the number of publications, while Total Link Strength (TLS) reflects the strength of collaboration among entities.

## 3. Results

### 3.1. Annual Scientific Publications and Cumulative Citation analysis

The number of publications per year serves as a fundamental bibliometric metric, offering valuable insights into the growth, productivity, and impact of research [19]. The findings of this study reveal that the number of annual scientific publications and cumulative citations in the field of gene editing and aging have shown an upward trend, which can be roughly divided into three phases. Phase I, from 2015 to 2016, had fewer than 50 publications per year. Phase II, from 2017 to 2019, saw a steady increase in publications, rising from 50 to over 100 per year. Phase III, from 2020 to 2024, experienced a slight decrease in publications in 2023 and 2024. The annual percentage growth rate of scientific publications was 21.08%. Bibliometric analysis in the gene editing and aging fields indicates that research activity and attention in this area have been consistently growing (Figure 1A).

### 3.2. Countries/Regions Contribution Analysis

#### 3.2.1. Country scientific production and Corresponding’s Countries

The findings of this study reveal the following distribution of articles across countries: USA: The USA has the highest number of articles (285), accounting for 29.17% of the total, with 23.16% of these being multinational collaborative papers. While it leads in absolute research volume, it has the lowest international collaboration rate. China: China ranks second with 214 articles, representing 21.90% of the total, and 24.30% of these are multinational collaborative papers. It is highly productive, with a slightly higher international collaboration rate than the USA. United Kingdom: The UK has fewer articles (51) but a significantly higher percentage of multinational collaborative papers (47.06%). Japan: Japan has 42 articles, with a relatively low percentage of multinational collaborative papers (14.29%). Germany: Germany has 41 articles, but a high percentage of multinational collaborative papers (48.78%). South Korea: South Korea has 37 articles, with a low percentage of multinational collaborative papers (8.11%). Canada: Canada has 29 articles, with 34.48% of them being multinational collaborative papers. Ethiopia: Ethiopia has 26 articles, with a low percentage of multinational collaborative papers (7.69%). Australia: Australia has 25 articles, with 40.00% of these being multinational collaborative papers. Italy: Italy has 18 articles, with 33.33% of them being multinational collaborative papers. The USA and China are the two countries with the highest number of articles, accounting for 29.17% and 21.90% of the total, respectively. The United Kingdom and Germany exhibit a high percentage of multinational collaborative papers, with 47.06% and 48.78%, respectively. Japan and South Korea have a lower proportion of multinational collaborative papers, with 14.29% and 8.11%, respectively (Figure 2B, 2C).

**Figure. 2A.**
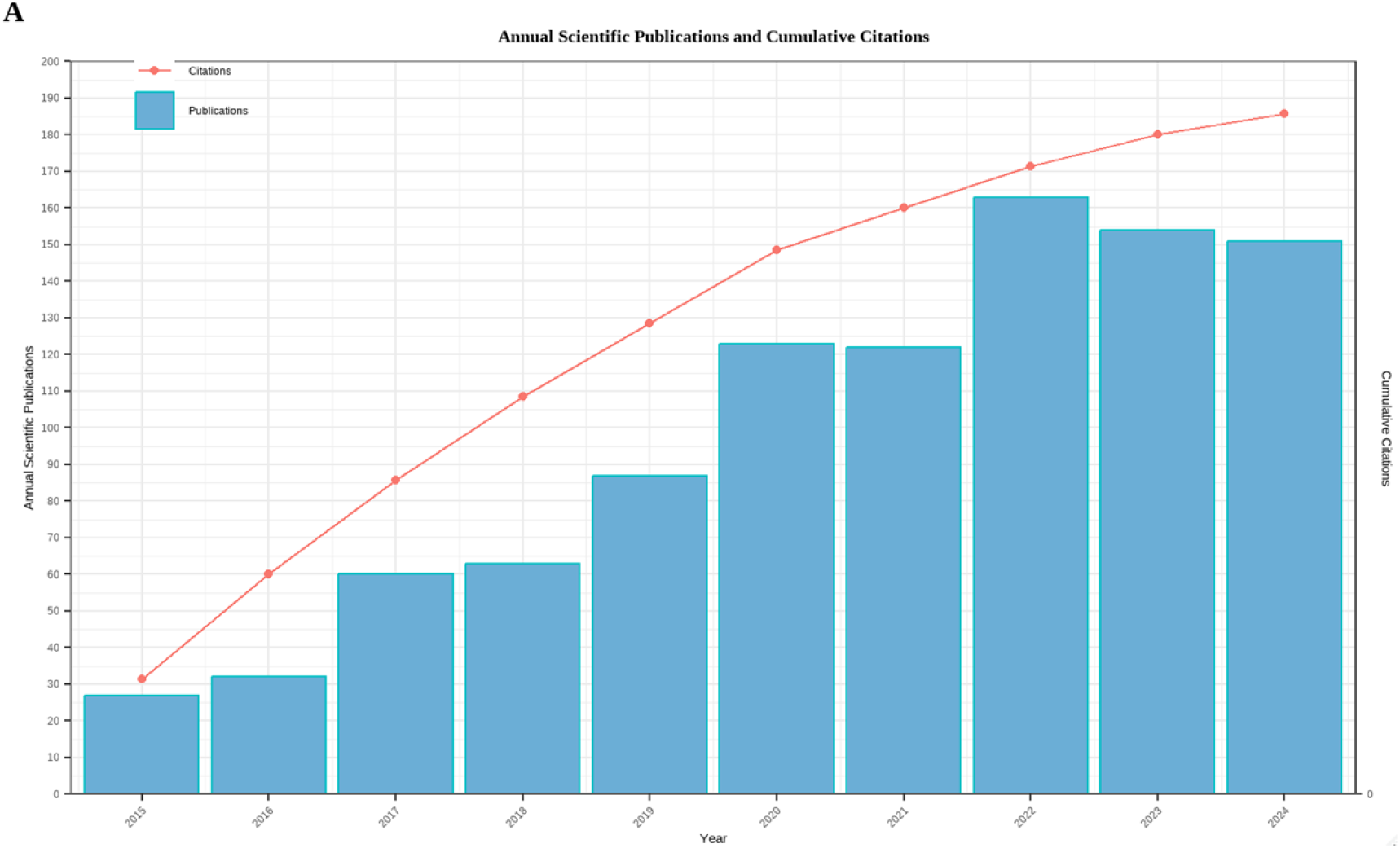
Annual Scientific Publications and Cumulative Citation from 2015 to 2024

**Figure. 2B.**
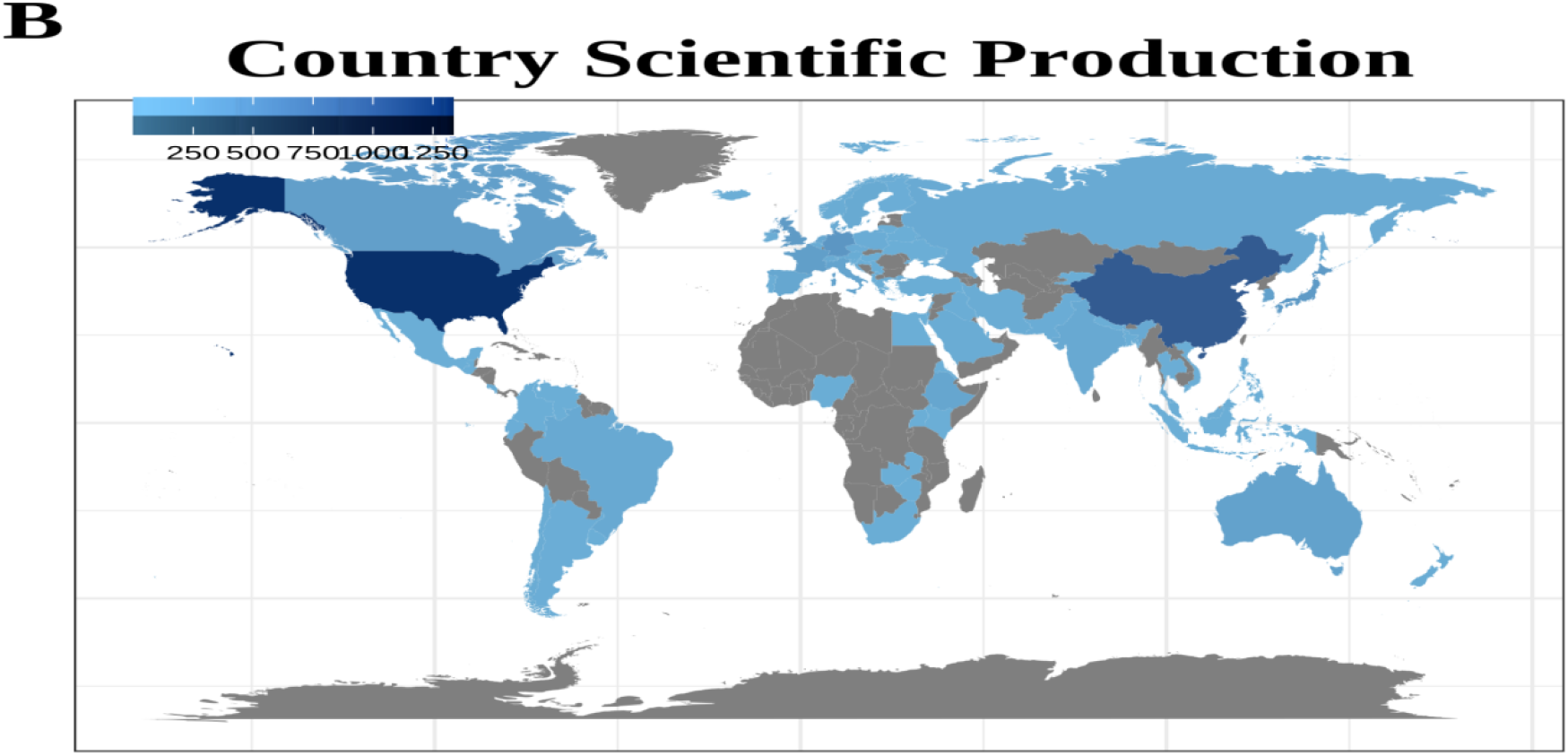
Global distribution of periodical articles published (2015-2024)

**Figure. 2C.**
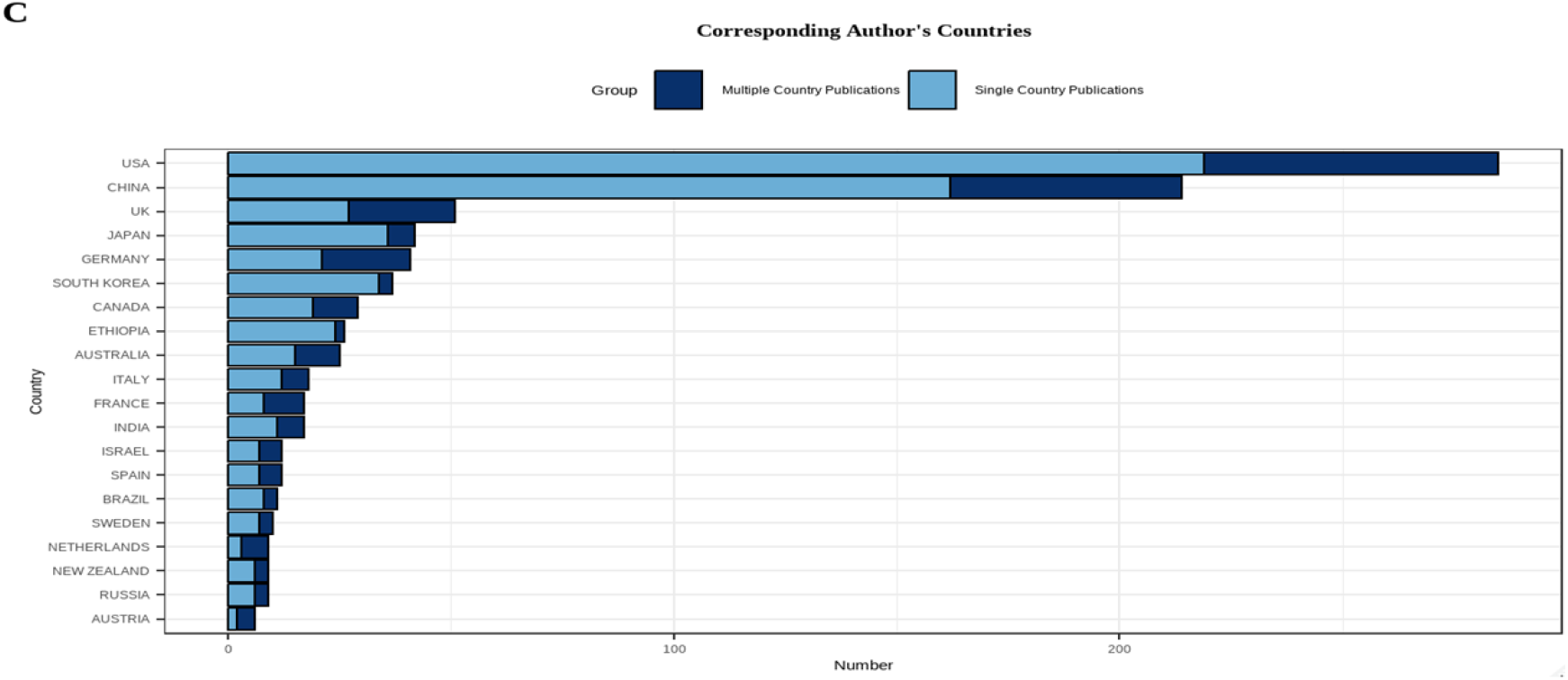
Corresponding Author’s countries

#### 3.2.2. Primary Interacting Countries and Frequencies

Our analysis of the Web of Science Core Database reveals that 53 countries/regions have published research on gene editing applications in aging. As shown in Figure 3, the top 30 international collaborative networks have formed through countries’ publications in this specific field. High-frequency interaction countries: The United States has the highest interaction frequency with other countries, particularly with China, Canada, France, Germany, and the United Kingdom. China maintains broad interactions with multiple countries, especially the U.S., the U.K., Germany, and Australia. The U.K., Germany, France, and Canada also frequently interact with both the U.S. and China. Asian countries such as China, Japan, and South Korea demonstrate close intra-regional collaboration. Medium-frequency interaction countries: Australia, Japan, the Netherlands, Italy, Spain, and Sweden engage in moderate-frequency interactions with multiple partners, making them key contributors to international research collaboration. Low-frequency interaction countries: Argentina, Brazil, Chile, Kenya, Malaysia, and Singapore, despite their relatively low interaction frequencies, are actively participating in international collaboration. African nations, notably Ethiopia and Kenya, are emerging on the global scientific stage, forging new research partnerships. Southeast Asian countries, particularly Malaysia and Singapore, are increasing their involvement in global research collaborations.

**Figure. 3.**
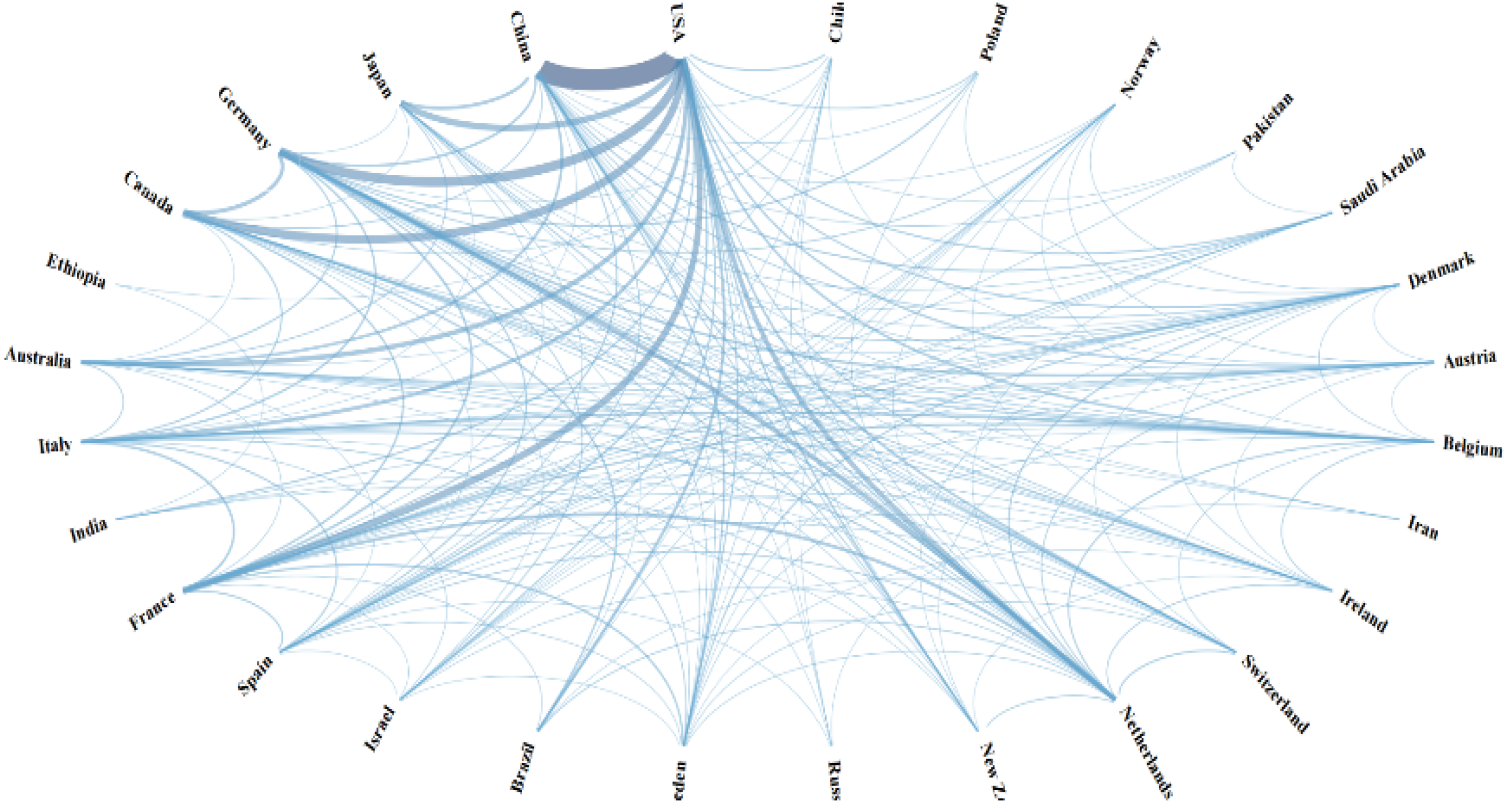
The cooperation network of the top 30 productive countries/territories. the width of the link is positively correlated with the strength of cooperation

### 3.3. Institutions contribution analysis

The top 15 productive institutions are shown in Figure 4. The majority of the most influential institutions are based in China and the USA, with North American institutions dominating the field of gene editing and aging research. Harvard University is a global leader in gene editing research, with 116 publications, closely followed by the University of California System with 90 publications. Harvard Medical School and its affiliated institutions contribute a total of 64 publications, with Harvard Medical School alone accounting for 49 publications, collectively establishing a first-tier research cluster and showcasing the region’s unparalleled expertise in this field. Asian institutions are also making significant strides. The Chinese Academy of Sciences, with 84 publications, Fudan University with 37 publications, and Seoul National University with 37 publications, all rank within the global top 10. In Europe, University College London (UCL), with 48 publications, stands out as a leader in aging and gene editing research, ranking among the top British institutions. The most influential institutions are located in China and the United States, driven by advancements in high-throughput sequencing technologies. Research in gene editing and aging exhibits a dual pattern: high concentration in the U.S. and China, coupled with global diffusion. Figure 5 clearly illustrates that the majority of publications in the WoS Core collection database come from universities and colleges (59%), followed by other agencies (18%), research institutes (13%), hospitals (7%), and clinical organizations (3%).

**Figure. 4.**
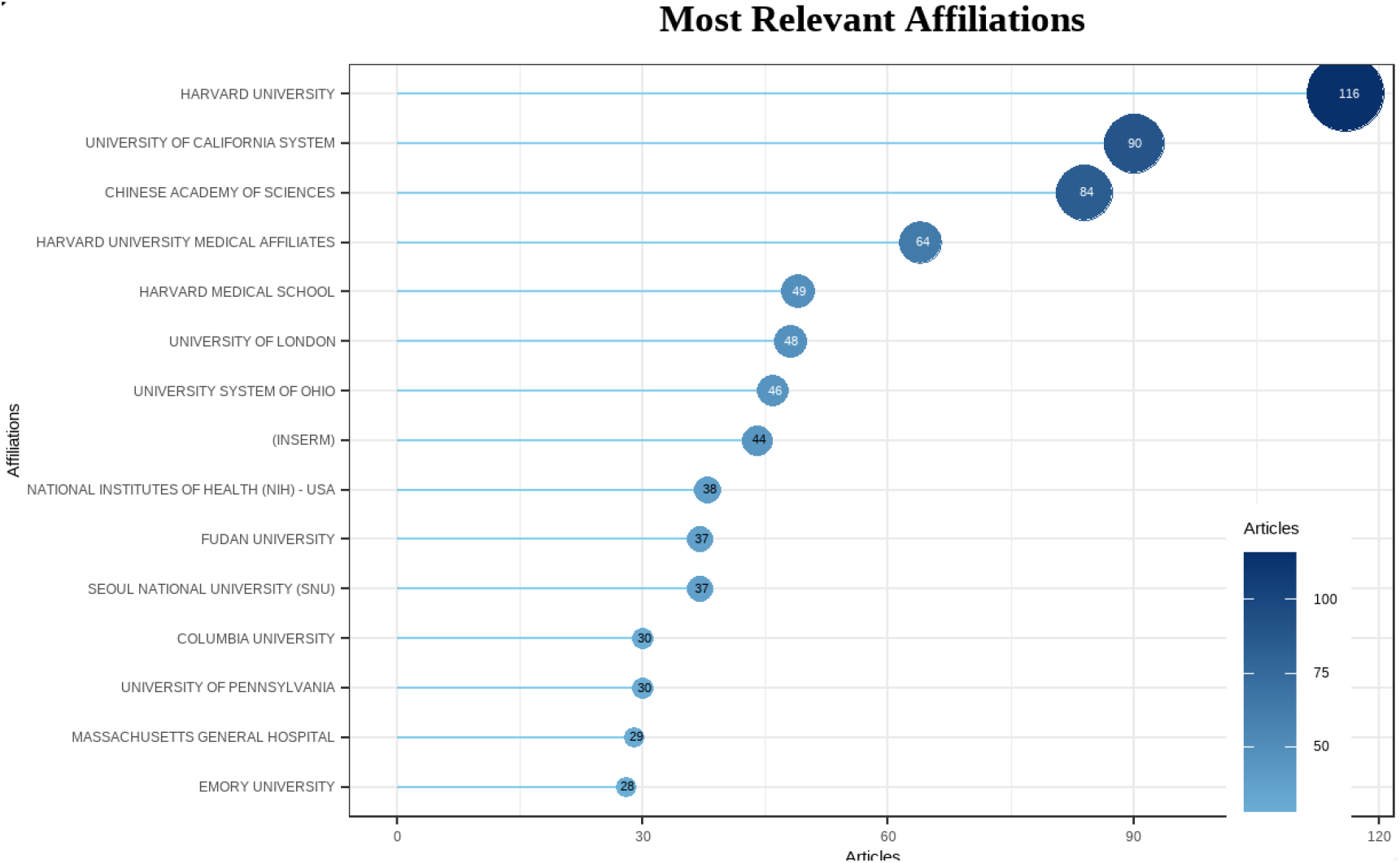
Top 15 most relevant affiliations

**Figure. 5.**
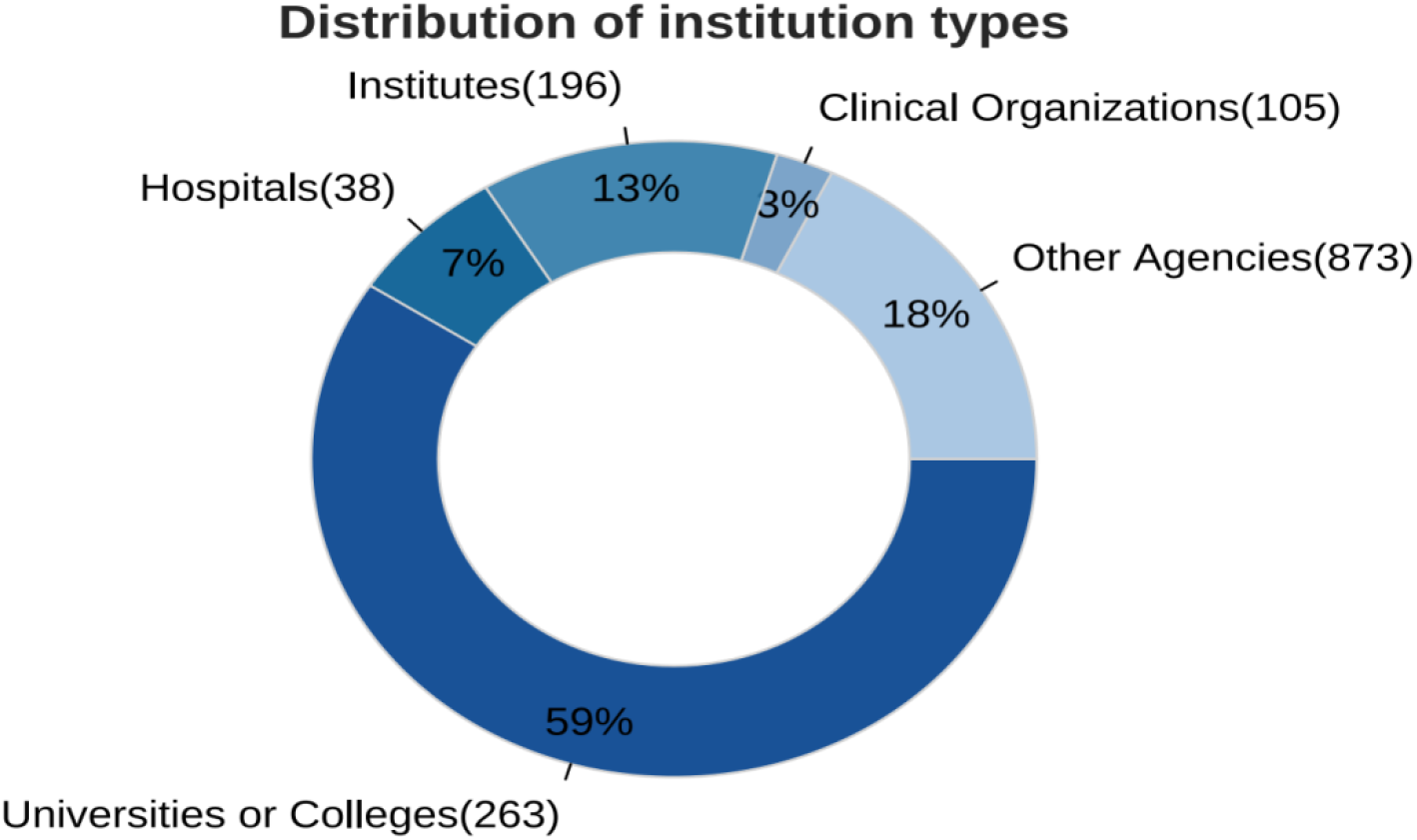
The distribution of institution types

### 3.4. journals analysis

As demonstrated in Figure 6, the number of publications in the five listed academic journals has shown consistent annual growth since 2015. SCIENCE REPORTS: There has been steady year-on-year growth since 2015, reaching 23 articles in 2024, making it one of the most prolific journals. INTERNATIONAL JOURNAL OF MOLECULAR SCIENCES: No publications were recorded until 2018. However, publications began in 2019 and experienced rapid annual growth, reaching 22 articles in 2024. PLOS ONE: This journal has experienced continuous growth since 2015, albeit at a decelerated rate after 2020, but still maintaining an upward trajectory. NATURE COMMUNICATIONS: Initially, this journal had low publication numbers, but growth accelerated after 2018, reaching 15 articles in 2024. FRONTIERS IN PLANT SCIENCE: Publications began in 2017, with an initial low output. However, growth surged post-2020, reaching 13 articles in 2024. Projections for 2025: It is clear that publications in the listed journals will continue to grow. As research deepens, these journals will become more specialized, creating more targeted platforms. Increased interdisciplinary studies and collaborations are expected to drive innovation and development in related fields.

**Figure. 6.**
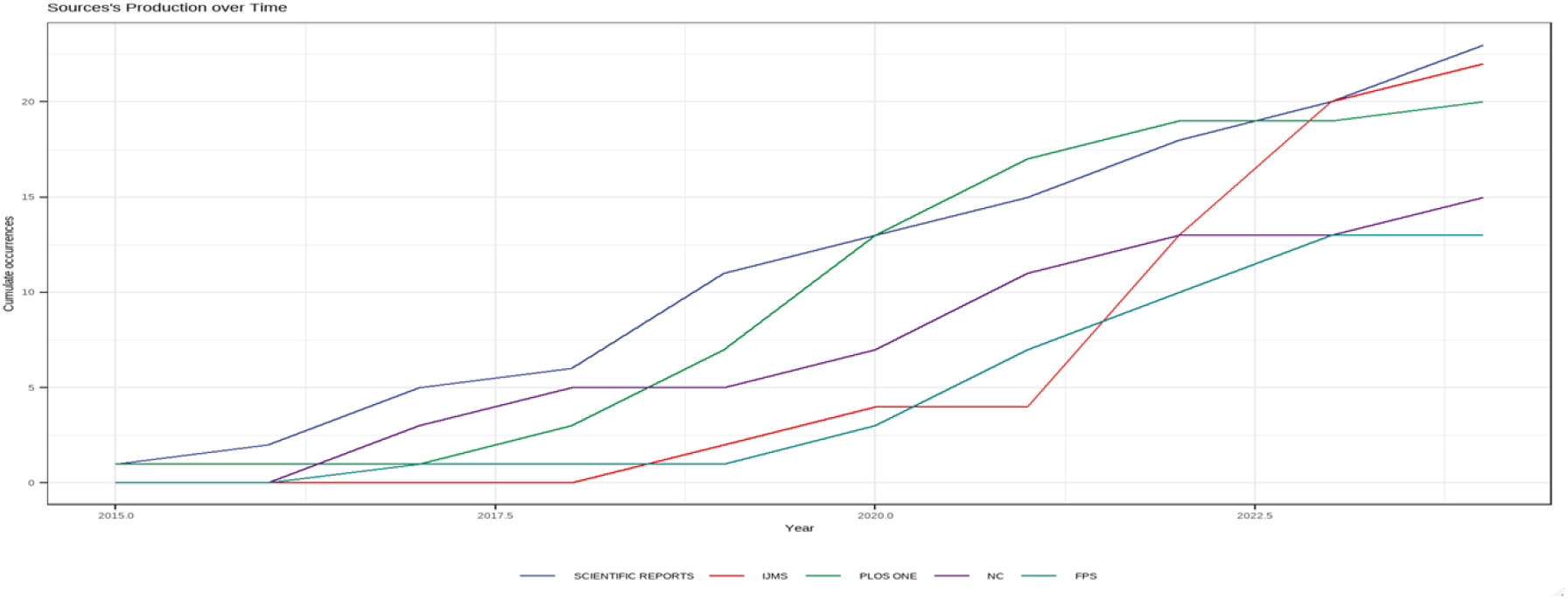
Annual growth curve of the top 5 institutions in the total number. IJMS: INTERNATIONAL JOURNAL OF MOLECULAR SCIENCES. NC: NATURE COMMUNICATIONS. FPS: FRONTIERS IN PLANT SCIENCE.

### 3.5. Authors and article analysis

The bibliometric analysis reveals the most prominent authors in the fields of gene editing and aging research, with their influence assessed based on publication output and citation impact. As demonstrated in Table 1, ZHANG Y leads in publication volume with 12 papers, though with a moderate citation impact (102 citations, H-index: 6). In contrast, KIM JH, with 11 publications, emerges as the most highly cited author (1,438 citations, H-index: 8), demonstrating significant influence in the field. Notably, KIM JH shows a strong collaborative relationship with KIM JS, as evidenced by their high co-citation count (702), suggesting a highly productive research partnership. LI Y also demonstrates significant impact, with 307 total citations and an average of 38.4 citations per paper, indicating consistent research relevance. This author’s collaboration with LIU GH (257 co-citations) further underscores the importance of strategic partnerships in enhancing citation influence. Additionally, KIM J (256 citations, avg. 32 citations/paper) and WANG Y (240 citations, avg. 24 citations/paper) contribute prominently to the field, reinforcing their roles as key researchers. These findings identify KIM JH and LI Y as central figures in gene editing and aging research, with collaborative networks playing a pivotal role in enhancing their scholarly impact. Furthermore, this study reveals that the average number of authors per article is 8.1, reflecting a common pattern of large collaborative teams. Only 61 (6.2%) single-authored articles were found, suggesting that the field of gene editing and aging is highly collaborative. The international coauthorship rate stands at 28.72%, which is considered an intermediate to high level, although lower than that in top disciplines like physics, where it often exceeds 50%.

**Table 1.** Top 10 authors and co-cited authors related to gene editing and aging.

| Rank | Author | Documents | Citations | Average-citations | H-index | Co-cited-author | Co-citations |
| --- | --- | --- | --- | --- | --- | --- | --- |
| 1 | ZHANGY | 12 | 102 | 8.50000 | 6 | MIAOY | 27 |
| 2 | KIMJH | 11 | 1438 | 130.7272 | 8 | KIMJS | 702 |
| 3 | WANGY | 10 | 240 | 24.00000 | 5 | LIUZ | 157 |
| 4 | LIUY | 9 | 108 | 12.00000 | 5 | ALFONSOJM | 40 |
| 5 | KIMJ | 8 | 256 | 32.00000 | 5 | RODIGSJ | 108 |
| 6 | LEEJ | 8 | 104 | 13.00000 | 4 | KIMDH | 45 |
| 7 | LIY | 8 | 307 | 38.37500 | 5 | LIUGH | 257 |
| 8 | ZHANGH | 8 | 122 | 15.25000 | 5 | WANGY | 59 |
| 9 | BERRYDP | 7 | 127 | 18.14286 | 3 | WALLE | 64 |
| 10 | CHENJ | 7 | 95 | 13.57143 | 5 | ADAMSONC | 32 |

As demonstrated in Table 2, KIM E (2017, NAT COMMUN): TC (Total Citations): 494, TC per Year: 54.9, NTC (Normalized Total Citations): 9.33. This paper, titled In vivo genome editing with a small Cas9 orthologue derived from Campylobacter jejuni, has the highest total citations in the field of gene editing, indicating a very high impact. It also has a high average number of citations per year, demonstrating its ongoing influence in the field. STROUD DA (2016, NATURE): TC: 385, TC per Year: 38.5, NTC: 6.46. This paper, Accessory subunits are integral for assembly and function of human mitochondrial complex I, also boasts a high citation count in gene editing and related fields. SHEN YJ (2022, IEEE T PATTERN ANAL): TC: 315, TC per Year: 78.8, NTC: 31.69. This paper has the highest average number of citations per year, despite its relatively recent publication, suggesting rapid accumulation of influence in the field of pattern analysis. KOBLAN LW (2021, NATURE): TC: 283, TC per Year: 56.6, NTC: 16.13. Published in 2021, this paper has an exceptionally high average number of citations per year, indicating it has garnered significant academic attention in a short period. MCDANIEL BT (2018, CHILD DEV): TC: 269, TC per Year: 33.6, NTC: 7.13. This paper has made an impact in the field of child development, suggesting that gene editing techniques are receiving growing attention for their potential applications in developmental biology. These papers cover the application of gene editing across various fields, including basic biology, disease mechanisms, developmental biology, pathogen research, stem cell research, and molecular therapy. High-Impact Papers: The papers by KIM E and STROUD DA stand out in terms of total citations and average citations per year, reflecting their central position in the field of gene editing. The papers by SHEN YJ and KOBLAN LW, although published later, have very high citation rates per year, indicating that these research areas are rapidly developing. Furthermore, the application of gene editing technology in fields such as child development and eating disorders demonstrates its interdisciplinary potential.

**Table 2.**
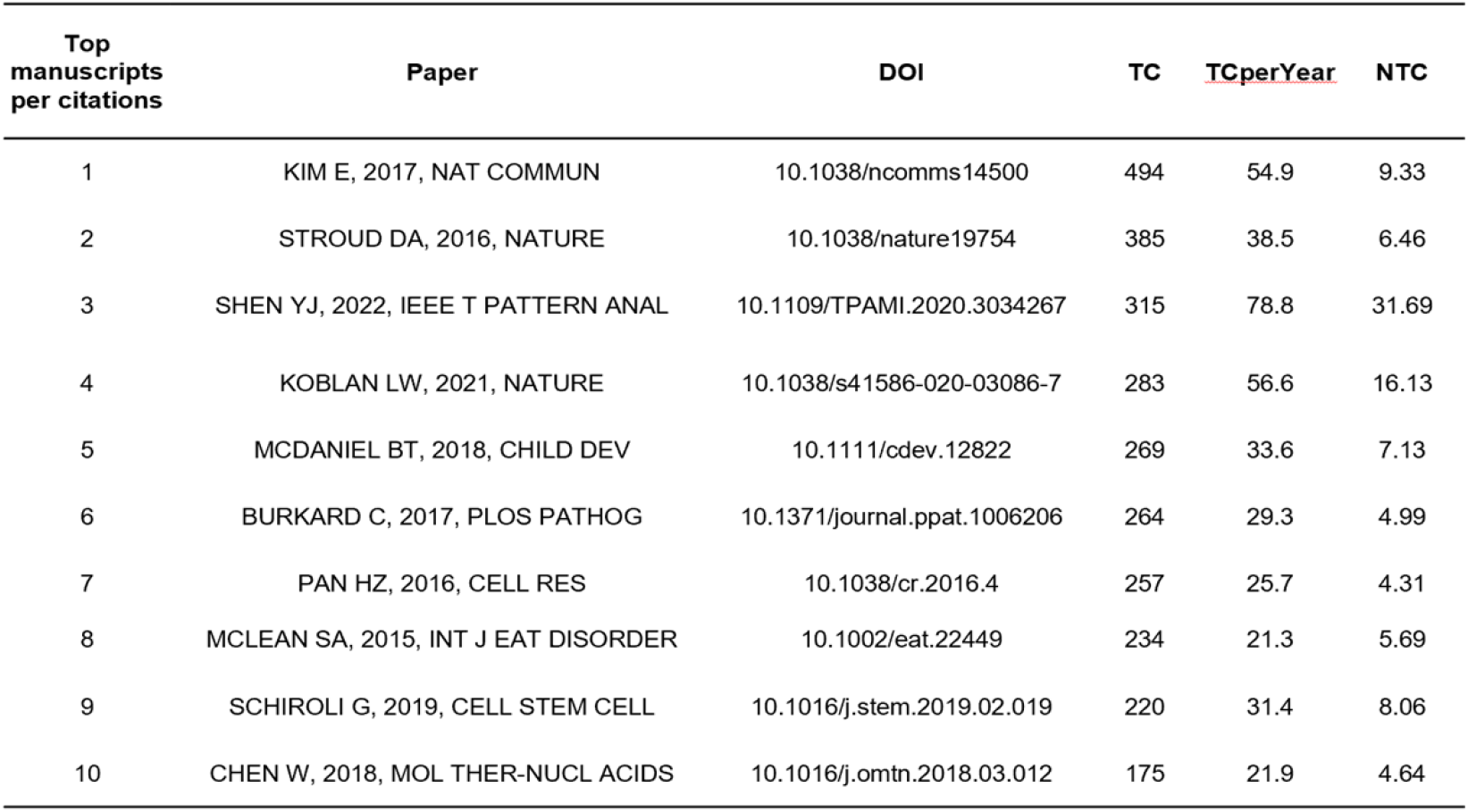
Top 10 papers ranked by citation count related to gene editing and aging.

| Top manuscripts per citations | Paper | DOI | TC | TCperYear | NTC |
| --- | --- | --- | --- | --- | --- |
| 1 | KIM E, 2017, NAT COMMUN | 10.1038/ncomms14500 | 494 | 54.9 | 9.33 |
| 2 | STROUD DA, 2016, NATURE | 10.1038/nature19754 | 385 | 38.5 | 6.46 |
| 3 | SHEN YJ, 2022, IEEE T PATTERN ANAL | 10.1109/TPAMI.2020.3034267 | 315 | 78.8 | 31.69 |
| 4 | KOBLAN LW, 2021, NATURE | 10.1038/s41586-020-03086-7 | 283 | 56.6 | 16.13 |
| 5 | MCDANIEL BT, 2018, CHILD DEV | 10.1111/cdev.12822 | 269 | 33.6 | 7.13 |
| 6 | BURKARD C, 2017, PLOS PATHOG | 10.1371/journal.ppat.1006206 | 264 | 29.3 | 4.99 |
| 7 | PAN HZ, 2016, CELL RES | 10.1038/cr.2016.4 | 257 | 25.7 | 4.31 |
| 8 | MCLEAN SA, 2015, INT J EAT DISORDER | 10.1002/eat.22449 | 234 | 21.3 | 5.69 |
| 9 | SCHIROLI G, 2019, CELL STEM CELL | 10.1016/j.stem.2019.02.019 | 220 | 31.4 | 8.06 |
| 10 | CHEN W, 2018, MOL THER-NUCL ACIDS | 10.1016/j.omtn.2018.03.012 | 175 | 21.9 | 4.64 |

### 3.6. keyword analysis

**VOSviewer** is a software tool designed for constructing and visualizing bibliometric networks [20]. Keywords that are closely linked within a network are typically clustered together. These clusters represent themes or topics that emerge from co-occurrence analysis. VOSviewer automatically identifies and labels these clusters based on the network structure, revealing which keywords are central to a field and highlighting the interconnections between different research areas [20]. We utilized keyword co-occurrence network analysis (using VOSviewer) to identify research hotspots, as well as shifting and emerging topics. By consolidating keywords with similar meanings, we allowed VOSviewer to seamlessly identify and label tightly associated keywords (Figure 7).

**Figure 7.**
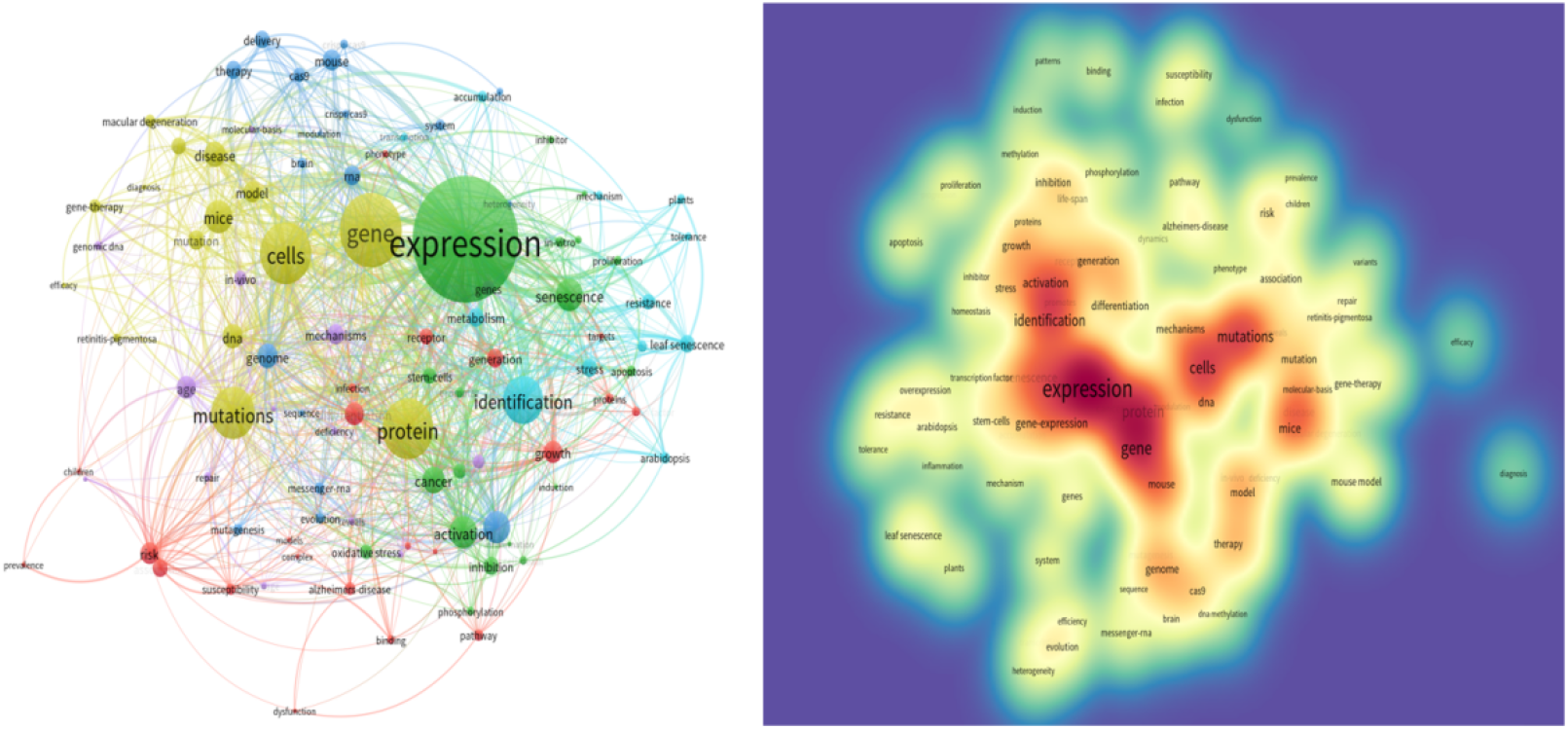
keyword co-occurrence network analysis. (A)The co-occurrence networks of 100 keywords were visualized by VOSviewer. (B)Large nodes represent keywords with relative higher occurrence; (C)Same color indicates relatively closer relationship; (D)All keywords could be grouped into four clusters.

#### 3.6.1. keyword co-occurrence network analysis

As demonstrated in Figure 7, the keywords can be categorized into four main clusters:Gene Expression Regulation (Red Cluster): This cluster includes keywords such as expression, gene-expression, messenger-RNA, and transcription factor. It primarily relates to mechanisms of gene expression regulation, including the roles of transcription factors and mRNA regulation.

Disease and Mutation (Green Cluster): Comprising keywords like mutations, risk, disease, gene-therapy, mouse, and mouse model, this cluster focuses on research into disease-related gene mutations, gene editing therapies (e.g., CRISPR-Cas9), and the use of animal models for studying genetic diseases.

Cellular and Molecular Mechanisms (Blue Cluster): Keywords in this cluster include cells, protein, DNA, mechanisms, gene, mouse, and model. It highlights research on protein-DNA interactions, cellular signaling pathways, and the functional validation of genes.

Plant Biology (Yellow Cluster): This cluster includes keywords such as plants, arabidopsis, and leaf senescence. It is centered on plant gene regulation, mechanisms of leaf senescence, plant resistance, and the function of senescence-related genes.

#### 3.6.2. Keyword co-occurrence dynamics Analysis

Keyword co-occurrence dynamics refer to the changes and evolving patterns in the co-occurrence of keywords or terms within a body of literature over time. This type of analysis enables researchers to trace how the relationships between concepts, topics, or terms shift in response to developments in research, technology, or societal needs [20]. Figure 8 illustrates the top 25 keyword co-occurrence dynamics from 2015 to 2024. **Early-Stage Hotspots (2015-2017)**: Keywords such as human cells, one-step generation, polymorphisms, and reliability peaked around 2016. These terms reflect foundational research in cell biology, validation of gene editing techniques, and studies of genetic diversity.

**Figure 8.**
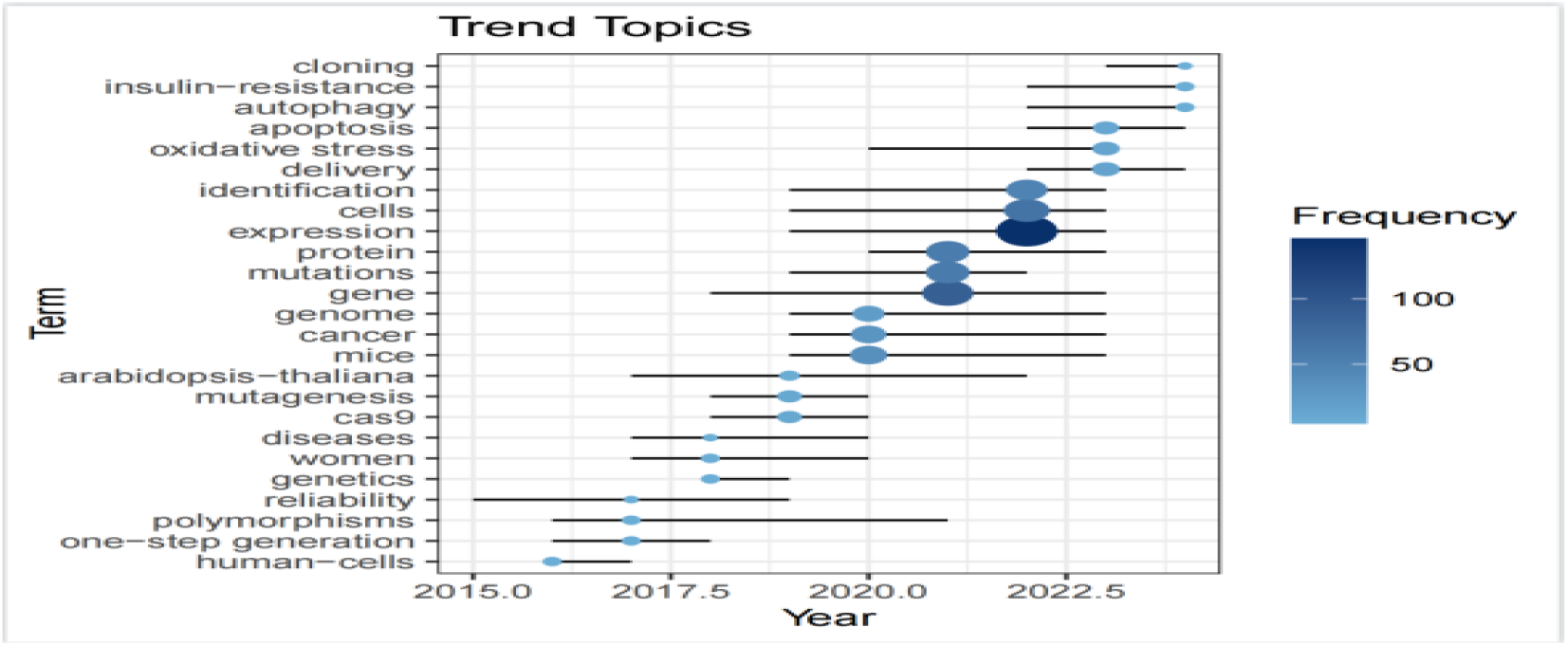
The TOP 25 Keyword co-occurrence dynamics related to gene editing and aging **4**.

**Mid-Term Hotspots (2018-2020)**: Terms like Cas9 and mutagenesis became dominant after 2018, indicating the rapid maturation of CRISPR-Cas9 technologies. Keywords such as mice, cancer, and genome (median year: 2019) signal a growing focus on animal models, cancer research, and genomic studies. Additionally, increased attention to women and diseases during this period suggests expanding clinical relevance and inclusivity in biomedical research.

**Recent Hotspots (2020-2024)**: More recent trends show rising interest in protein expression, cells, and identification, signaling advances in proteomics, cellular mechanisms, and gene function validation. Emerging terms like mutations and delivery systems (post-2020) highlight a deepening focus on off-target effects and gene delivery strategies. Since 2022, keywords such as oxidative stress, apoptosis, and autophagy have seen significant growth, reflecting intensified research into cell death and stress response pathways. Clinical innovations are also becoming prominent, with insulin resistance and cloning indicating increasing interest in metabolic disorders and cloning technologies.

## Discussion

In the era of big data, the pace of information generation is accelerating. Bibliometric analysis (BA) which focuses primarily on academic productivity utilizes published scientific literature (e.g., research articles, books, conference proceedings) to quantitatively assess research activity within a specific domain [21]. BA serves as a vital tool for rapidly mapping the landscape of emerging scientific fields. For the first time, this study conducts a comprehensive bibliometric analysis of literature related to Gene Editing and Aging. It systematically examines and reveals patterns across countries, institutions, authors, journals, publication trends, and keyword distributions within this research area. This analysis not only identifies the most prolific contributors and influential institutions but also provides valuable insights for funding agencies and policy makers in allocating resources more effectively toward researchers and organizations that consistently produce high-quality work. Furthermore, the findings can assist scholars in identifying potential collaborators and uncovering promising subfields for further exploration.

### 4.1. Current Status and Major Contributing Countries

Our analysis of the Web of Science core database shows that 53 countries/regions have published research on gene editing applications in aging. The United States leads globally with 285 articles. Regional clusters indicate that Europe/North America (USA, Germany, UK) and East Asia (China, Japan, Korea) dominate the field. The extensive scientific exchanges between countries, along with the increasing number of cross-regional partnerships and the active participation of emerging economies, highlight the breadth and inclusiveness of global scientific collaboration. Research activity and interest in this area continue to grow, with both the number of annual publications and cumulative citations steadily increasing.

### 4.2. Active Institutions and Authors

Researchers often analyze authors’ affiliations to determine which universities or research institutions are most active in a particular field. This helps to understand the research output of different institutions. This study shows that Harvard University ranks first globally with 116 publications, followed by the University of California system (90 publications) and the Chinese Academy of Sciences (84 publications) in third place. An analysis of contributions by institutional affiliation shows that universities/colleges have the largest share (59%), followed by other institutions (18%), research institutes (13%), hospitals (7%), and clinical organizations (3%).

The h-index is a single numerical indicator that summarizes the research impact of authors. A unique feature of the h-index is that it takes into account both the number of publications and their citation impact, providing a balanced measure of an individual’s or institution’s research output [17]. KIM JH is the most influential author with 11 papers, 1,438 citations (130.7 average citations), and an h-index of 8. His partnership with KIM JS is particularly prominent (702 total co-citations). ZHANG Y has the highest number of publications (12), making him the most prolific author, though with a moderate citation impact (102 total citations, h-index of 6). LI Y has an average of 38.4 citations per article (307 total citations), demonstrating outstanding research continuity and a significant collaborative network with LIU GH (257 total co-citations), KIM J (32 citations per article), and WANG Y (24 citations per article). The average number of authors per article is 8.1, and only 6.2% of the articles are single-authored, indicating a strong reliance on teamwork. The international collaboration rate is 28.72%, reflecting a moderately high level of globalization and cooperation. The study is highly dependent on collaboration, and core authors have enhanced their academic impact through high-quality partnerships, particularly the collaboration involving KIM JH.

### 4.3. Active Journals

The analysed journals have demonstrated sustained growth in publications since 2015, with Scientific Reports and the International Journal of Molecular Sciences emerging as the most productive. Projections for 2025 indicate a continuation of this growth trajectory, driven by increasing research specialisation and interdisciplinary collaboration, which are likely to foster further innovation in the field.

### 4.4. Research Hotspots and Frontier Trend

In bibliometrics, keyword co-occurrence analysis and co-cited references are used to visualize the knowledge network and frontier research in a field [22].This study uses VOSviewer’s high-frequency word statistics, co-occurrence network, and temporal sequence evolution methods to systematically identify hot directions in gene editing and aging: prioritizing CRISPR optimization (e.g., KOBLAN LW study) and AI-assisted analysis (SHEN YJ study). It also considers the potential translational value of plant gene research (e.g., leaf senescence).Keyword co-occurrence networks and sequential evolution analysis reveal a dynamic progression in gene editing and aging research, characterized by three distinct phases: Tool development (2015-2017);Model validation (2018-2020); Clinical translation (2021-present).

#### 4.4.1. Tool Development (2015-2017)

In the early phase of this study, the keywords “human cells,” “one-step generation,” “polymorphisms,” and “reliability” peaked in 2016, reflecting foundational research in cell biology, gene editing validation, and genetic diversity. The time-dependent accumulation of cellular damage is widely considered the general cause of aging [23]. Genome integrity and stability are pervasively threatened by exogenous chemical, physical, and biological agents, as well as by endogenous challenges such as DNA replication errors, chromosome segregation defects, oxidative processes, and spontaneous hydrolytic reactions. The wide range of genetic lesions caused by these extrinsic or intrinsic sources of damage includes point mutations, deletions, translocations, telomere shortening, single- and double-strand breaks, chromosomal rearrangements, defects in nuclear architecture, and gene disruption caused by the integration of viruses or transposons. All these molecular alterations and the resulting genomic mosaicism may contribute to both normal and pathological aging [24]. Other forms of damage, such as chromosomal aneuploidy and copy-number variations, are also associated with aging. These DNA alterations may affect essential genes and transcriptional pathways, resulting in dysfunctional cells that may ultimately compromise tissue and organismal homeostasis. This is especially relevant when DNA damage affects stem cells, hampering their role in tissue renewal or leading to their exhaustion, which in turn promotes aging and increases susceptibility to age-related pathologies [11, 25]. Moreover, studies in humans and other long-lived species have shown that enhanced DNA repair mechanisms coevolve with increased longevity [26].

Since the advent of human gene sequencing technology in 1977 [12]], humanity has progressed from gene sequencing to gene editing, which refers to a technology that enables precise modification of DNA sequences at the genomic level. This is achieved by using sequence-specific nucleases (e.g., ZFN, TALEN, and CRISPR/Cas9) to create DSBs in the DNA, thereby inducing cellular repair mechanisms such as NHEJ or HDR to facilitate targeted gene insertion, deletion, or replacement [12, 16, 27]. In 2012, two research groups published findings stating that purified Cas9, derived from Streptococcus thermophilus or Streptococcus pyogenes, can be guided by crRNAs to cleave target DNA in vitro [27].

#### 4.4.2. Model Validation (2018-2020)

By 2018, the dominant keywords shifted to “Cas9” and “mutagenesis,” marking the maturation of CRISPR/Cas9 technology. CRISPR/Cas9 is a simple and rapid tool that enables efficient modification of endogenous genes in various species and cell types. Moreover, a modified version of the CRISPR/Cas9 system with transcriptional repressors or activators allows robust transcription repression or activation of target genes. The simplicity of CRISPR/Cas9 has resulted in the widespread use of this technology in many fields, including basic research, biotechnology, and biomedicine [15]. By relying on sgRNA to guide the Cas9 nuclease for targeted DNA cleavage, this approach offers high efficiency, low cost, and multiplex editing capabilities, making it the current mainstream technology. In 2019, keywords predominantly centered on “mice” and “cancer,” highlighting research priorities in animal models, oncology, and genomics. Concurrently, there was an increased focus on women and diseases, reflecting an expansion in clinical relevance. In this study, the evolution of keywords follows the research trajectory of “tool development → model validation → clinical translation.” At the same time, the growth of PLOS ONE article publications slows down after 2020, and the frequency and reliability of mutations are negatively correlated with a peak in 2023, suggesting that the safety challenges of gene editing continue to escalate and that safer editing tools may be needed in the future [28].

#### 4.4.3. Clinical Translation (2021-present)

The 2021 keyword “cloning renaissance” may be related to the breakthrough of gene-edited cloned animals and the ensuing ethical debate. Keyword correlation analysis predicts that, in the short term (1-2 years), the link between autophagy and metabolic disease may drive breakthroughs in precision medicine. In the long term (3-5 years), the convergence of CRISPR-Cas9/synthetic biology (cloning) could redefine biotechnology, prioritizing dual validation (technical + clinical), such as Cas9 + cancer. It is important to avoid isolated clusters (e.g., cloning). Meanwhile, increasing mutations and decreasing reliability may trigger public distrust as a risk warning. The high frequency of the keyword delivery and the long-term low frequency of genetics after 2020 may hinder the development of this technology. Delivery in the field of gene editing is the key technical link to realize gene modification tools (e.g., CRISPR/Cas9 systems, base editors, etc.) and accurately reach the target cells or tissues. Harnessing CRISPR-Cas9 technology for cancer therapeutics has been hampered by low editing efficiency in tumors and potential toxicity of existing delivery systems [29, 30]. Despite the extensive utilization of gene editing technologies such as CRISPR/Cas9 in non-human primate models, their suboptimal knock-in efficiency and off-target effects continue to compromise the precision and reliability of target screening. For instance, CRISPR/Cas9 technology exhibits low gene editing efficiency in non-human primate embryos and carries potential off-target risks [31]. In the present study, Daniel Rosenblum et al. describe a safe and efficient lipid nanoparticle (LNP) that uses a novel aminoconjugated lipid for the delivery of Cas9 mRNA and sgRNA [30]. Gene editing technology, a revolutionary tool in the life sciences, has demonstrated considerable potential in the treatment of diseases. However, it has also given rise to unprecedented ethical controversies, which extend beyond the domains of scientific safety and technological boundaries, encompassing profound issues such as the nature of human life, social justice, and intergenerational responsibility.

#### 4.4.4. Future Perspectives

The future, guided by gRNA, will enable precise DNA targeting. Subsequent advancements in base editing (BE) and prime editing (PE) have enabled single-base substitutions or short insertions without the need for double-strand breaks [32, 33]. Current core challenges include improving target design efficiency, controlling off-target effects, and developing new tools. The explosive growth of AI has revolutionized these aspects through data-driven modeling and generative design, propelling gene editing towards intelligent, high-efficiency evolution. The integration of AI with gene editing, particularly in off-target effect prediction, protein design, and delivery optimization [34, 35], signals the transition of the life sciences from “describing nature” to “programming life.” This transformation accelerates scientific discovery and reshapes future medical paradigms. As CRISPR co-discoverer Jennifer Doudna stated, “AI gives us the power to redesign life, but we must use it responsibly.” To ensure equitable global benefit, we must balance technological innovation with ethical governance. In the future, it is expected that people will achieve a leap in the quality of their lives and extend their healthy lifespan through multidisciplinary cross-innovation and global collaboration.

## 5. Limitations

This study revealed several limitations due to the inherent nature of any bibliometric analysis. First, since the database is periodically updated, there may be incomplete literature retrieval, which could introduce selection bias. Second, we selected only the Web of Science Core Collection database as our data source. Incorporating other databases, such as PubMed,Scopus, and Google Scholar, would increase the robustness of our findings. Finally, this study exclusively focused on English literature, while excluding non-English publications. Moreover, this study did not analyze or compare the funding sources of different studies.

## 6. Conclusion

We summarized the publication information of gene editing and aging-related literature from 2015 to 2024, including data on countries, institutions, authors, journals, and keywords. We then analyzed the research hotspots based on these publications and predicted future research frontiers. This first bibliometric analysis of gene editing and aging reveals rapid growth since 2015, led by the U.S., China, and Europe. The field has progressed through three phases: tool development (2015-2017), model validation (2018-2020), and clinical translation (2021-present). Future directions in this field include CRISPR optimization and AI-assisted analysis. Additionally, the potential translational value of plant gene research (e.g., leaf senescence) should be considered. Current core challenges include improving target design efficiency, controlling off-target effects, and developing new tools.

## Supplementary material

来自网页 : (TS=((Editing, Gene) OR (Genome Editing)OR (Editing, Genome)OR (Base Editing)OR (Editing, Base))) AND TS=( (Biological Aging) OR (Aging, Biological)OR (Aging)OR(Senescence)) – 1,739 – Web of Science Core Collection

## Notes

### Competing Interest Statement

The authors have declared no competing interest.

